# Ferroptotic pores induce Ca^2+^ fluxes and ESCRT-III activation to modulate cell death kinetics

**DOI:** 10.1101/867564

**Authors:** Lohans Pedrera, Rafael A. Espiritu, Uris Ros, Anja Schmitt, Stephan Hailfinger, Ana J. García-Sáez

**Affiliations:** Institute for Genetics and Cologne Excellence Cluster on Cellular Stress Responses in Aging-Associated Diseases (CECAD), Köln-Universität, Joseph-Stelzmann-strasse 26, 50931 Köln, Germany; Interfaculty Institute of Biochemistry, Eberhard-Karls-Universität Tübingen, 72076 Tübingen, Germany

**Author notes:** Equal contribution (Lohans Pedrera, Rafael A. Espiritu). Department of Chemistry, De La Salle University, 2401 Taft Avenue, Manila 0922, Philippines (Rafael A. Espiritu).

## Abstract

Ferroptosis is an iron-dependent form of regulated necrosis associated with lipid peroxidation. Despite its key role in the inflammatory outcome of ferroptosis, little is known about the molecular events leading to the disruption of the plasma membrane during this type of cell death. Here we show that a sustained increase in cytosolic Ca^2+^ is a hallmark of ferroptosis that precedes complete bursting of the cell. We report that plasma membrane damage leading to ferroptosis is associated with membrane nanopores of few nanometers in radius and that ferroptosis, but not lipid peroxidation, can be delayed by osmoprotectants. Importantly, Ca^2+^ fluxes during ferroptosis correlate with the activation of ESCRT-III-mediated membrane repair, which counterbalances the kinetics of cell death and modulates the inflammatory signature of ferroptosis. Our findings with ferroptosis provide a unifying concept that sustained high levels of cytosolic Ca^2+^ prior to plasma membrane disruption are a common feature of regulated necrosis and position ESCRT-III as a general protective mechanism in these inflammatory cell death pathways.

## Introduction

Ferroptosis is a caspase-independent form of regulated cell death characterized by the generation of lethal amounts of iron-dependent lipid hydroperoxides in cellular membranes (Dixon, Lemberg et al., 2012, Yang & Stockwell, 2016). Cells dying via ferroptosis are characterized by plasma membrane rupture and release of otherwise confined intracellular components including pro-inflammatory damage-associated molecular patterns (DAMPs) (Proneth & Conrad, 2019). Therefore, this type of cell death is associated with necroinflammation and with activation of the innate immune system. In the context of tissue injury and degeneration, ferroptosis has been shown to contribute to diseases related to ischemia/reperfusion injury (Li, Feng et al., 2019, Tuo, Lei et al., 2017), tissue damage and organ demise (Conrad, Angeli et al., 2016), and to a number of pathologies including neurodegenerative diseases and cancer (Stockwell, Angeli et al., 2017). For these reasons, understanding the molecular mechanisms involved in ferroptosis is not only of biological, but also of medical relevance.

In contrast to other cell death pathways that possess a specific cellular machinery to mediate cell death, ferroptosis seems to be executed by engaging cellular components of the metabolic machinery as a response to failure of antioxidant mechanisms (Dixon, 2017). Polyunsaturated fatty acid metabolism and the generation of specific lipid peroxides through iron-dependent enzymatic reactions are key processes in this form of cell death (Agmon & Stockwell, 2017, Yang, Kim et al., 2016). The best characterized trigger of ferroptosis is depletion or inhibition of Glutathione Peroxidase 4 (GPX4), a unique enzyme responsible for reducing peroxidized phospholipids in membranes to their less toxic alcohol equivalents (Seiler, Schneider et al., 2008, Yang et al., 2016). Treatment with Ras-selective Lethal small molecule 3 (RSL3), which directly inhibits GPX4, or with erastin-1, which affects the catalytic cycle of GPX4 by diminishing the levels of its co-substrate glutathione (GSH), are broadly used methods to induce ferroptosis (Dixon et al., 2012, Dixon, Patel et al., 2014, Yang, SriRamaratnam et al., 2014). In addition to the canonical glutathione-based GPX4 pathway, a recent study showed that the ferroptosis supresor protein 1 (FSP1) targets ubiquinone in the plasma membrane and reduces it to ubiquinol in order to protect cells from lipid peroxidation and ferroptotic cell death (Bersuker, Hendricks et al., 2019, Doll, Freitas et al., 2019)

A key step in ferroptosis execution is the final disruption of the plasma membrane, which mediates the release of intracellular factors acting as DAMPs. However, the molecular mechanism causing the loss of plasma membrane integrity during ferroptosis remains completely unsettled and the nature and size of the membrane injury in ferroptosis has remained unexplored. Damage to the plasma membrane in other forms of necrotic death like necroptosis (Cai, Jitkaew et al., 2014, Ousingsawat, Cabrita et al., 2017, Ros, Pena-Blanco et al., 2017), pyroptosis (de Vasconcelos, Van Opdenbosch et al., 2018, Ruhl, Shkarina et al., 2018), or upon chemical, toxin, or laser-induced cell death (Espiritu, Pedrera et al., 2019) results in the activation of ion fluxes. In these scenarios, Ca^2+^ fluxes resulting from membrane damage have been linked with the activation of membrane repair mechanisms (Andrews & Corrotte, 2018, Etxaniz, Gonzalez-Bullon et al., 2018, Jimenez & Perez, 2017). The Endosomal sorting complexes required for transport (ESCRT) machinery seems to play a critical counterbalancing role that can delay or even block cell death in necroptosis and pyroptosis (Gong, Guy et al., 2017a, Gong, Guy et al., 2017b, Ruhl et al., 2018).

Here we investigated the molecular mechanism of plasma membrane permeabilization during ferroptosis triggered by erastin-1 and RSL3 in mouse fibroblasts. Using live-cell imaging and flow cytometry, we defined and tracked in parallel the kinetics of different hallmarks of ferroptotic cell death. We show that ferroptosis progression involves production of peroxidized lipids prior to a sustained increase of cytosolic Ca^2+^ and final plasma membrane breakdown. We also identify the formation of nanopores as a core mechanism that triggers plasma membrane burst during ferroptosis, which consequently can be inhibited by osmoprotectants of proper size. Finally, we correlate the increase in cytosolic Ca^2+^ in ferroptosis with the activation of the ESCRT-III machinery, which acts as a protective mechanism that delays plasma membrane disruption and cell death. This has physiological consequences, as depletion of ESCRT-III modulates the secretion of the anti-inflammatory cytokine IL-10 in ferroptosis, thereby reshaping the microenvironment and the inflammatory signature of ferroptotic cells. Our findings support a general role of the ESCRT-III machinery in counterbalancing membrane damage and in modulating the inflammatory outcome during regulated necrosis.

## Results

### Sustained increase in cytosolic Ca^2+^ is a hallmark of ferroptosis

Similar to other types of regulated necrosis such as necroptosis and pyroptosis, ferroptosis leads to complete plasma membrane disruption and the final bursting of the cell (Dixon et al., 2012, Proneth & Conrad, 2019). Increase in cytosolic Ca^2+^, cell swelling, rounding, as well as plasma membrane breakdown have been described as hallmarks for necroptosis and pyroptosis, which represent critical differences to apoptosis (Frank & Vince, 2018, Ros et al., 2017, Vandenabeele, Galluzzi et al., 2010). To assess whether these cellular alterations also take place during ferroptosis, we measured cytosolic Ca^2+^ levels in parallel to changes in cell morphology and plasma membrane breakdown by live-cell confocal imaging at 37 °C (Figure 1A and B) and by flow cytometry (Figure 1C-H) in mouse fibroblasts (NIH-3T3 cells) treated with the ferroptosis inducers erastin-1 or RSL3. We used the non-fluorescent marker fluo-4 acetoxymethyl (Fluo-4 AM), which is cleaved inside cells to yield the impermeant, fluorescent form of the Ca^2+^ indicator (Stosiek, Garaschuk et al., 2003), to visualize alterations in the levels of cytosolic Ca^2+^ and the fluorescent DNA-intercalating agent propidium iodide (PI) as marker of irreversible plasma membrane disruption and cell death (Rieger, Nelson et al., 2011). In parallel, we quantified changes in cell shape from transmission microscopy images using Image J (Ros et al 2017).

**Figure 1:**
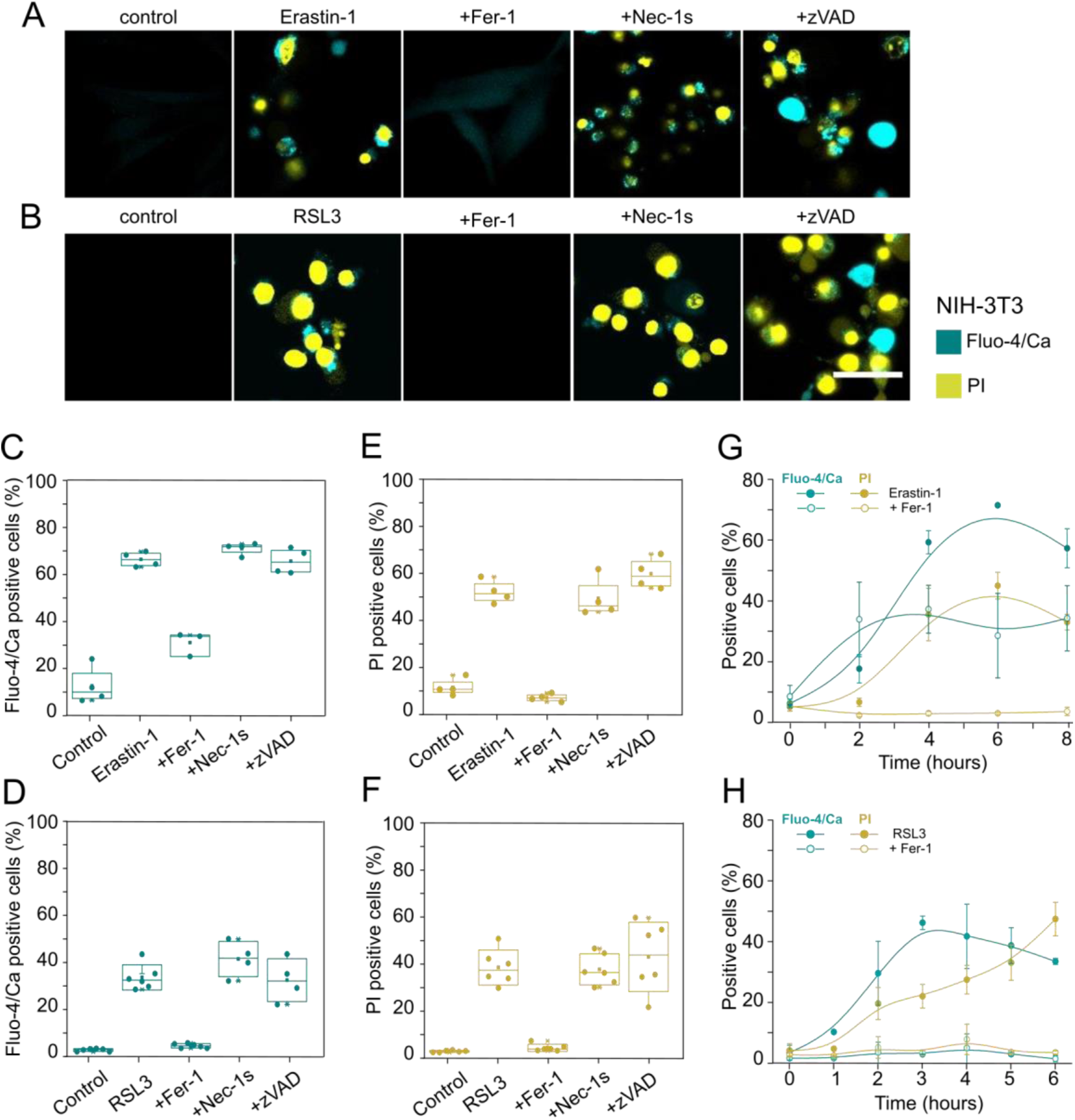
Levels of cytosolic Ca^2+^ increase during ferroptosis in a treatment-dependent manner. A-B) Confocal images of NIH-3T3 cells treated with Erastin-1 (A) or RSL3 (B) in the presence or not of different cell death inhibitors. Pictures are representative of at least three independent experiments. Scale bar, 50 µm. C-F) Counting by flow cytometry of Fluo4-AM and PI positive cells treated with erastin-1 (C and E) or RSL3 (D and F) in the presence or not of different cell death inhibitors. G-H) Kinetics of increase of cytosolic Ca^2+^ and plasma membrane breakdown in cells treated with Erastin-1 (G) or RSL3 (H) in the presence or not of Fer-1. The values represent the mean and the standard deviation of at least three independent experiments. Increase in cytosolic Ca^2+^ signal was detected using the Ca^2+^ indicator Fluo4-AM and plasma membrane breakdown was detected with PI. Fer-1: ferroptosis inhibitor ferrostatin 1, Nec-1s: necroptosis inhibitor necrostatin-1s, zVAD: pan-caspase inhibitor for apoptosis and pyroptosis. Concentrations: Erastin-1 (10 µM), RSL3 (2 µM), Fer-1 (2 µM), Nec-1s (10 µM), zVAD (20 µM).

We detected a clear increase in cytosolic Ca^2+^ concentration in the cytosol of NIH-3T3 cells upon treatment with erastin-1 or with RSL3 (Figure 1A and B). This was accompanied by cell shape changes including the appearance of a single swelling bleb followed by complete plasma membrane breakdown or bursting. Similar events were visualized when human fibrosarcoma (HT-1080) or human breast carcinoma cells (Mda-157) were treated with RSL3 (Figure S1). This sequence of morphological alterations was similar to what we previously found during necroptosis (Ros et al., 2017), but less dramatic than during toxin-induced necrosis (Ros et al., 2017) or pyroptosis (Chen, He et al., 2016, Frank & Vince, 2018), in which most of the cells remain attached to the surface upon swelling. Increase in cytosolic Ca^2+^, cell rounding (as a proxy for cell shape changes) and plasma membrane breakdown were specifically inhibited in the presence of the ferroptosis inhibitor ferrostatin 1 (Fer-1) (Dixon et al., 2012), but not by the necroptosis inhibitor necrostatin-1s (Nec-1s) nor the pan-caspase inhibitor zVAD (Gong et al., 2017b, Ros et al., 2017), suggesting that these events are specific hallmarks of ferroptosis (Figure 1A-F). Similar to NIH-3T3 cells, Fer-1 also completely inhibited the increase in cytosolic Ca^2+^ and cell death in HT-1080 and Mda-157 cells upon treatment with RSL3 (Figure S1).

To further characterize the temporal relationship between cytosolic Ca^2+^ increase and plasma membrane disruption upon erastin-1 and RSL3 treatment in the NIH-3T3 population, we tracked in parallel the kinetics of Fluo-4/Ca^2+^ and PI fluorescence by flow cytometry (Figure 1G and H). Independently of treatment, increase in cytosolic Ca^2+^ was an early event that took place hours before plasma membrane breakdown. We also found that changes in cytosolic Ca^2+^ levels were not completely abolished by Fer-1 in cells treated with erastin-1, with about 40% of the cell population remaining Fluo-4/Ca^2+^ positive (Figure 1C and G). In contrast, Fer-1 completely hindered cytosolic Ca^2+^ increase when ferroptosis was induced with RSL3 (Figure 1D and H). These results suggest the activation of two distinct Ca^2+^ signaling processes by erastin-1: one that is unrelated with ferroptosis, and a later one, specific to ferroptosis. This late increase in cytosolic Ca^2+^ can be considered a hallmark of ferroptosis, as it was inhibited by Fer-1 and was common for erastin-1 and RSL3-induced ferroptosis.

### Lipid oxidation precedes increase in cytosolic Ca^2+^, and plasma membrane breakdown

Our results above suggested that Ca^2+^ signaling is an early event during ferroptosis compared to plasma membrane disruption. However, the detailed temporal relationship between Ca^2+^ increase and other ferroptosis hallmarks in bulk cell experiments is masked by the heterogeneity in which the cells in the population undergo ferroptosis upon treatment. To address this issue, we quantified the increase in Fluo-4/Ca^2+^ signal, cell rounding, and PI intake at the single cell level using live cell confocal microcopy. We took cell rounding and detachment as an internal reference to compare the relative kinetics of ferroptosis hallmarks between individual cells in independent experiments (Figure 2). From the kinetic curves obtained from single cells (Figure 2C, F and J), we calculated the time required to achieve 50% change of each ferroptotic phenotype (t_50_). This allowed us to set the relative lag time between each process in response to ferroptosis induction with erastin-1 or RSL3 (Figure 2D, G, and K).

**Figure 2:**
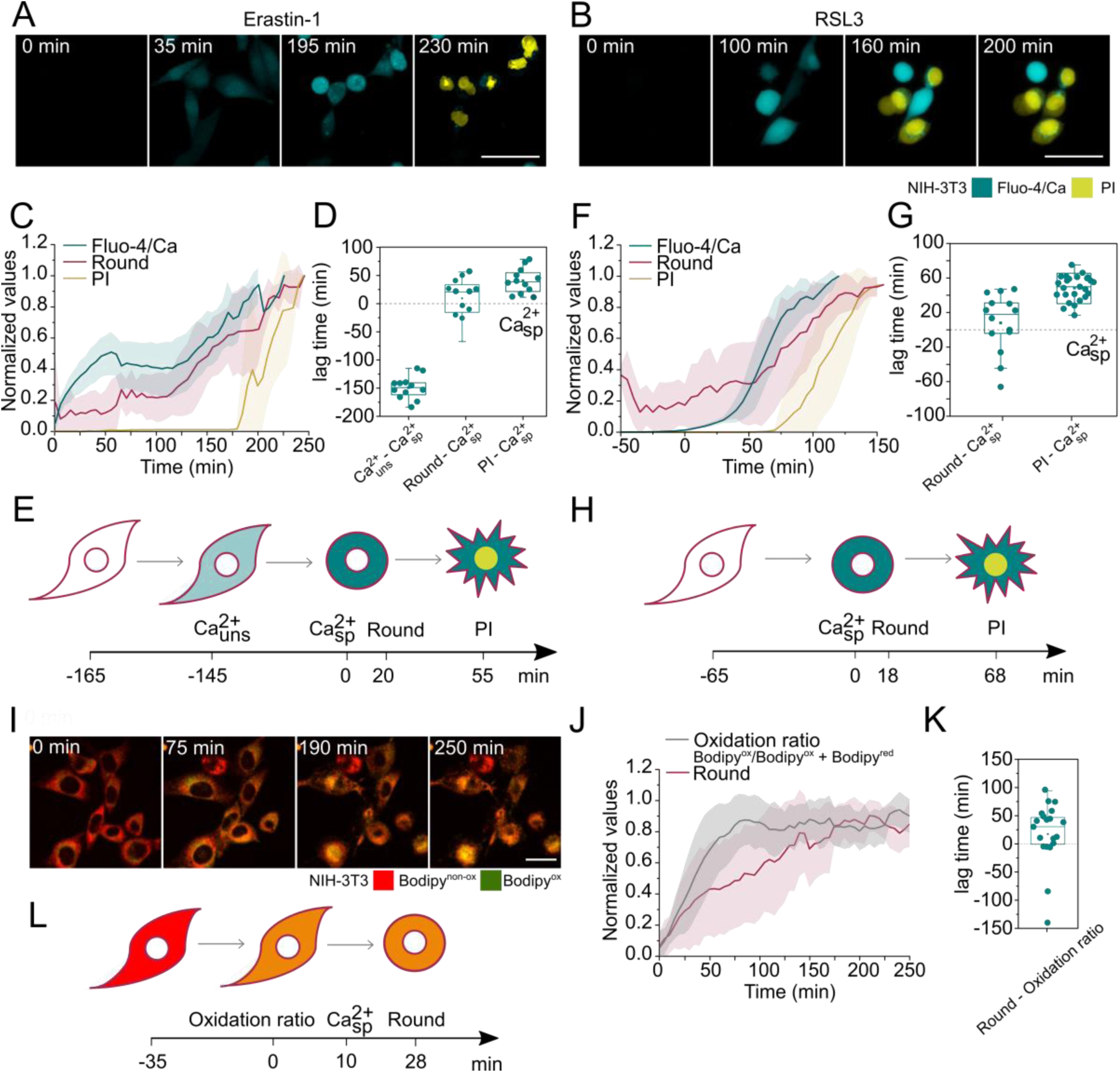
Specific increase in cytosolic Ca^2+^ takes places prior to plasma membrane breakdown. A-B) Representative time-lapse images of NIH-3T3 cells treated with (A) Erastin-1 or (B) RSL3, and monitored for cell rounding and Fluo-4AM and PI staining. Scale bar, 50 µm. C) Kinetics of increase of the normalized Ca^2+^ signal, change in shape, and PI intake upon erastin-1 treatment. Plots show the average (N = 16 cells) temporal relationships between the normalized parameters. All cells were synchronized to the first appearance of specific Ca^2+^ signal (*t* = 0). D) Time delay between specific Ca^2+^ flux and other ferroptotic events (unspecific Ca^2+^, change in shape and PI) observed upon Erastin-1 treatment. E) Graphical representation of the sequence of events observed during Erastin-1-induced ferroptosis. F) Kinetics of increase of the normalized Ca^2+^ signal, change in shape and PI intake upon RSL3 treatment. Plots show the average (N = 25 cells) temporal relationships between the normalized parameters. All cells were synchronized to the first appearance of specific Ca^2+^ signal (*t* = 0). G) Time delay between specific Ca^2+^ flux and other ferroptotic events (change in shape and PI) observed upon RSL3 treatment. H) Graphical representation of the sequence of events observed during RSL3-induced ferroptosis. I) Representative time-lapse images of lipid peroxidation in NIH-3T3 cells upon treatment with RSL3. Scale bar, 25 µm J) Kinetics of increase of the normalized Bodip ratio and change in shape upon RSL3 treatment. Plots show the average (N = 21 cells) temporal relationships between the normalized parameters. All cells were synchronized to the increase in lipid peroxidation ratio signal (*t* = 0). K) Time delay between cell round and lipid peroxidation ratio observed upon RSL3 treatment. L) Graphical representation of the sequence of the lag time observed between lipid peroxidation and other ferroptotic events during RSL3-induced ferroptosis. In C, F and J, values were normalized between 0 and 1. In D, G and K t_50_ of each event was calculated from individual curves obtained per single cells (shown in C, F and J). These values correspond to the time at 50% of the maximum signal. Concentrations: Erastin-1 (10 µM) and RSL3 (2 µM).

Independently of the ferroptotic trigger, increase in cytosolic Ca^2+^ anticipated cell rounding and complete plasma membrane collapse (Figure 2A and B). However, as suggested by the flow cytometry experiments (Figure 1G), increase in cytosolic Ca^2+^ in the presence of erastin-1 was a two-step process with biphasic behavior (Figure 2C). The first event reached saturation around 40% of the maximum Ca^2+^ signal and corresponded to the unrelated calcium (Ca^2+^_un_) increase, as it was not observed with RSL3 (Figure 2F) and was not inhibited by Fer-1 (Figure 1G). The second Ca^2+^ rise, common to RSL3 treatment and inhibited by Fer-1 (Figure 1H and Figure 2F) corresponded to the cytosolic Ca^2+^ increase specific to ferroptosis (Ca^2+^_sp_). Temporally, Ca^2+^_un_ increased happened after erastin-1 treatment, followed by Ca^2+^_sp_ rise, cell rounding, and final PI intake (Figure 2C-E). In contrast, RSL3-treated cells were characterized by a unique and specific Ca^2+^ rise event, which took place later after treatment, followed by cell rounding and final PI intake (Figure 2F-H). Given the fact that RSL3 induced only the ferroptosis-specific Ca^2+^ signal, we selected this drug for further experiments aimed at characterizing the connection of this ion with lipid peroxidation and membrane damage in ferroptosis.

We next analyzed the relative kinetics of Ca^2+^ increase and lipid peroxidation at the single cell level. We tracked in parallel the change in the ratio between the red and green fluorescence signal of the lipid peroxidation sensor C11 BODIPY 581/591 and the cell rounding (Figure 2 I and J). RSL3 treatment promoted lipid peroxidation detected by the increase in the green fluorescent signal of BODIPY (BODIPY_ox_) (Figure 2I), which preceded the rounding of the cells (Figure 2J and K). Based on the interpolation of the relative kinetics of increase of the Fluo-4/Ca^2+^ (Figure 2F-H) and BODIPY_ox_ signals (Figure 2 J and K), we estimated that Ca^2+^ fluxes took place after lipid peroxidation during ferroptosis (Figure 2L).

### Osmotically active agents protect cells against ferroptosis

Pore formation is a common feature in the execution of different types of regulated cell death including apoptosis (Cosentino & Garcia-Saez, 2017), necroptosis (Ros et al., 2017) and pyroptosis (Sborgi, Ruhl et al., 2016). The opening of membrane pores leads to a net influx of water molecules and cell swelling as a consequence of the osmotic imbalance resulting from the high concentration of large intracellular molecules that cannot pass through membrane pores (Figure 3A). Such an effect can be prevented by addition to the external medium of osmotic protectants of appropriate size that are not able to enter the cell through the pores, thus counterbalancing the intracellular osmotic pressure, water influx, and the consequent cell collapse (Figure 3B) (Sukhorukov, Imes et al., 2009, Tejuca, Dalla Serra et al., 2001). According to this, the permeability of membrane pores, but not of ion channels, can be modulated by the osmotic effects exerted by polyethylene glycols (PEGs), which can be used to distinguish the nature of alterations in membrane permeability (Ros et al., 2017, Sborgi et al., 2016, Tejuca et al., 2001). To investigate whether the plasma membrane permeabilization observed in ferroptotic cells were connected with membrane pores, we assessed the effect of PEGs of different sizes on the kinetics and extent of different ferroptosis hallmarks.

**Figure 3:**
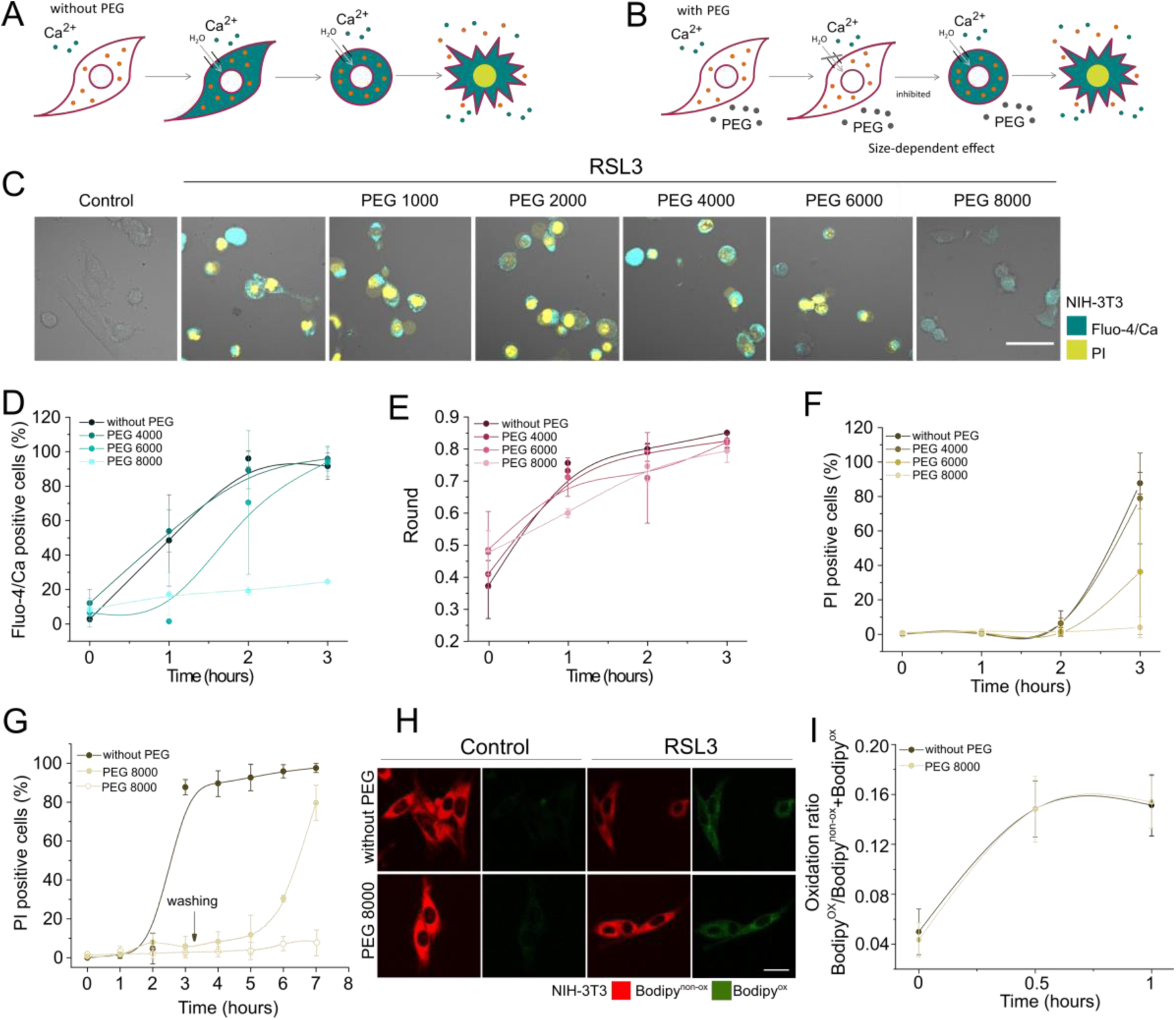
PEGs of high sizes provide osmotic protection against ferroptosis. A) Impermeant intracellular molecules impose an osmotic gradient after pore opening that leads to net influx of water molecules and cell lysis. B) PEGs of proper size can prevent this effect if their size is large enough to do not cross the membrane through the pores. C) Representative images of the effect of PEGs of different sizes on RSL3-induced ferroptosis at 3 hours of treatment. Scale bar, 50 µm. D) Kinetics of increase of cytosolic Ca^2+^, E) change in cell shape, and F) PI intake in NIH-3T3 cells treated with RSL3, in the presence or not of PEGs of different sizes. Each data point represents the mean of at least six replicas made in three independent experiments. At least 100 cells were analyzed for each replica. G) Recovery of cell death after washing out PEG 8000. NIH-3T3 cells were treated with RSL3 in the presence of PEG 8000 and after 3 hours the osmoprotectant was removed or not and the kinetics of cell death was tracked over time. H) Representative images of the effect of PEG 8000 in the oxidation ratio of C11 BODIPY 581/591 on RSL3-induced ferroptosis at 1 hours of treatment in NIH-3T3 cells. Scale bar, 25 µm. K) Oxidation ratio of C11 BODIPY 581/591 in cell treated with RSL3 in the presence or not of PEG 8000. Each data point represents the mean of at least 5 replica made in three independent experiments. At least 100 cells were analyzed for each condition. Concentrations: RSL3 (2 µM), PEGs (5 mM). PEG sizes: 4000 (1.8 nm), 6000 (2.3 nm) and 8000 (2.7 nm).

We used the following PEGs (in parenthesis their hydrated radii as reported in (Ling, Jiang et al., 2013, Tejuca et al., 2001)): 400 (0.56 nm), 600 (0.69 nm), 1000 (0.94 nm), 2000 (1.6 nm), 4000 (1.8 nm), 6000 (2.3 nm), and 8000 (2.7 nm). Addition of smaller PEGs up to 4000 did not prevent the increase of cytosolic Ca^2+^ signal, cell rounding or cell death (Figure 3C-G and S2). In contrast, both the increase in Ca^2+^ levels and cell death were delayed in the presence of higher-molecular weight PEGs (6000 and 8000) in a size dependent manner (Figure 3C-F and S2D). Complete protection of cell death and inhibition of Ca^2+^ fluxes were only achieved in the presence of PEG 8000. Under these conditions, we also observed a delay in the kinetics of cell rounding (Figure 3E). These results indicate that the increase in cytosolic Ca^2+^, cell rounding, and plasma membrane breakdown are all events driven by osmotic forces. Taking as a reference the size of the PEG able to protect, we could roughly estimate the range of the perforated sections of the plasma membrane to be around 2.3-2.7 nm radius. Notably, the protective effect of PEG 8000 in ferroptosis was reverted upon washing, which indicates that the membrane damage was stable over time (Figure 3G). We also found that PEG 8000 did not prevent lipid peroxidation in RSL3-treated NIH-3T3 cells (Figure 3H and I). These results confirmed that the effect of PEG 8000 was not a consequence of its ability to inhibit RSL3 or to block the ferroptotic pathway, but rather the result of counterbalancing the water and ion fluxes resulting from pore formation in the plasma membrane. Altogether, our data support a general model of cell disruption in ferroptosis through the formation of pores of a few nanometers radius in the plasma membrane.

### ESCRT-III complex is activated during ferroptosis and antagonizes cell death

The repair of plasma membrane damage induced by a variety of stimuli is an automatic response of the cell initiated by the acute and local increase in cytosolic Ca^2+^ concentration, as a consequence of greater membrane permeability (Andrews & Corrotte, 2018, Jimenez & Perez, 2017). It has been previously reported that during necroptosis and pyroptosis membrane repair dependent on the action of the ESCRT-III complex delays cell death (Gong et al., 2017a, Gong et al., 2017b, Ruhl et al., 2018).

To investigate the role of ESCRT-III complex in ferroptosis, we transiently transfected NIH-3T3 cells with the component CHMP4B, tagged with either eGFP or mCherry, and monitored its dynamics using live-cell confocal imaging. CHMP4B initially showed a homogenous cytosolic distribution (Gong et al., 2017b, Ruhl et al., 2018), however, upon ferroptosis induction with RSL3, the protein localized to distinct punctae that increased in number and fluorescence intensity over time (Figure 4A and B and S3). This observation was similar to the reported behavior of CHMP4B during necroptosis (Gong et al., 2017a, Gong et al., 2017b) and pyroptosis (Ruhl et al., 2018), and indicates that the ESCRT-III complex is likewise activated as part of the repair mechanisms during ferroptosis.

**Figure 4.**
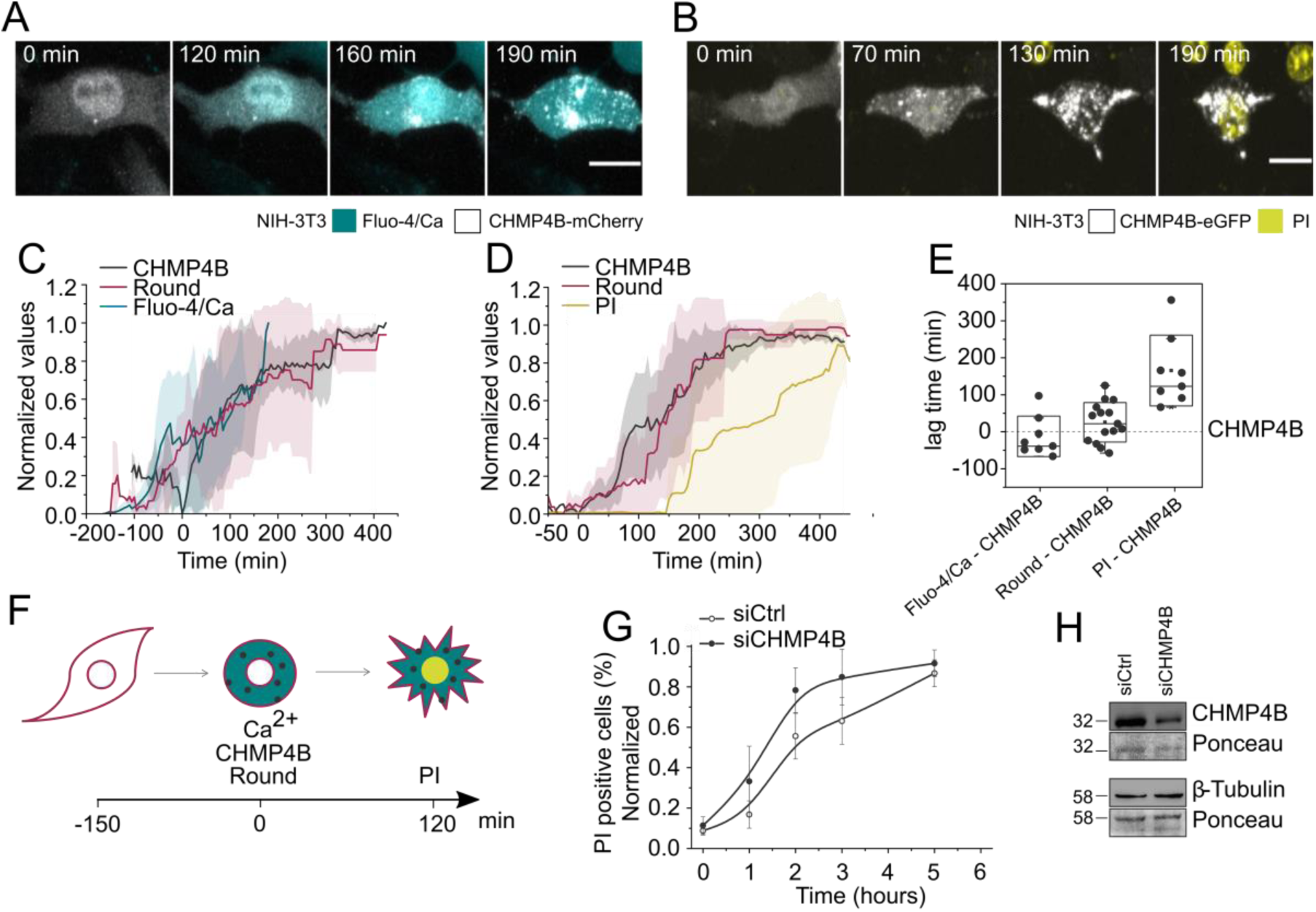
Membrane damage-induced Ca^2+^ entry correlates with CHMP4B activation to protect from cell death via membrane repair. A-B) Representative time-lapse images of NIH-3T3 cells transiently transfected with A) CHMP4B-mCherry or B) CHMP4B-eGFP, treated with RSL3, and monitored for CHMP4B punctae formation, Fluo-4 AM and PI staining. Scale bar, 20 µm. C-D) Kinetics of appearance of CHMP4B punctae and increase of cytosolic Ca^2+^ signal, change in shape, and PI intake upon RSL3 treatment. Plots show the average temporal relationships between normalized C) mCherry (N = 8 cells) or D) eGFP (N = 15 cells) fluorescence intensity standard deviation, cell rounding, Fluo-4 AM corrected total cell fluorescence (CTCF, calculated as integrated density – (Area x background intensity)), and PI fluorescence intensity. All cells were synchronized to the first appearance of CHMP4B punctae (*t* = 0). E) Time delay between appearance of CHMP4B punctae, Ca^2+^ signal, cell rounding and PI. t_50_ of each event was calculated from individual curves obtained per single cells (shown in C and D). These values correspond to the time at 50% of the maximum signal and were plotted as lag time with respect to appearance of CHMP4B punctae. F) Graphical representation of the sequence of events related to CHMP4B activation in ferroptosis. G) Effect of knocking down CHMP4B on the kinetics of ferroptosis. H) Immunoblot showing the knock down of CHMP4B. In G and H, NIH-3T3 cells were transfected with siRNA CHMP4B (20 µM) for 48 hours.

We next quantified the temporal correlation of CHMP4B punctae formation with the increase in cytosolic Ca^2+^ concentration, cell rounding, and final plasma membrane breakdown at the single cell level by live-cell confocal imaging. The start of CHMP4B punctae formation coincided approximately with the increase in cytosolic Ca^2+^ concentration (Figure 4C and E and S3). This is consistent with the fact that repair mechanisms are activated shortly after the rise in cytosolic Ca^2+^concentration (Andrews & Corrotte, 2018, Jimenez & Perez, 2017), and also with previous results showing formation of CHMP4B punctae after induction of membrane damage (Gong et al., 2017a, Gong et al., 2017b, Jimenez, Maiuri et al., 2014, Ruhl et al., 2018). Furthermore, the results showed that cells started to round up at about the same time of these two events (Figure 4C–E), consistent with a putative role of osmotic forces, as discussed in the previous section. Final membrane permeabilization occurred after sustained increase in cytosolic Ca^2+^ levels, cell rounding and ESCRT-III complex activation, likely when the cellular repair machineries were ultimately overwhelmed (Figure 4B, E and F).

To examine the functional role of ESCRT-III in ferroptosis, we assessed the effect of knocking down CHMP4B on the extent and kinetics of cell death in NIH-3T3 cells (Figure 4G and H). As shown in Figure 4G, we observed a higher and faster incidence of cell death in CHMP4B knock down cells compared with the control, particularly at early time points (1-3 hours), indicating that the loss of CHMP4B results in higher sensitization of the cells towards ferroptosis. Importantly, these findings show that the ESCRT-III machinery plays a role at protecting cells against ferroptosis most likely by counterbalancing membrane damage.

### ESCRT-III modulates the secretion of the anti-inflammatory cytokine IL-10 during ferroptosis

During necroptosis and pyroptosis, the ESCRT-III machinery has been shown to regulate the release of cytokines and other inflammatory factors, which affects the communication with immune cells (Gong et al., 2017a, Gong et al., 2017b, Ruhl et al., 2018, Yoon et al., 2017). We then examined the impact of ESCRT-III on cytokines secretion by assessing the effect of knocking down CHMP4B on the cytokine profile obtained from the supernatant of ferroptotic cells. We identified that IL-10 and IL-11 were specifically reduced in the supernatant of ferroptotic cells, which was reverted by depletion of CHMP4B (Figure 5A). Real time PCR quantification of transcriptional induction of IL-10 showed that the mRNA expression levels were kept during ferroptosis induction (Figure 5B). In stark contrast, knocking down of CHMP4B induced the upregulation of the *il-10* transcription in ferroptotic cells. Altogether, our results indicate that, similar to necroptosis and pyroptosis, ESCRT-III also modulates the cytokine profile of ferroptotic cells.

**Figure 5.**
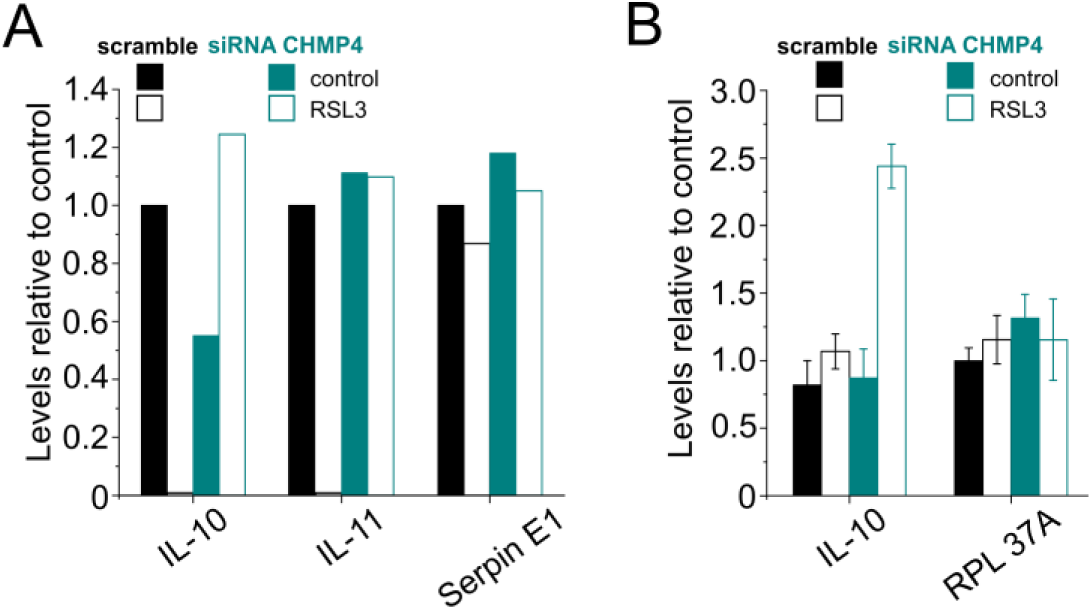
CHMP4B activation restricts the secretion of IL-10. A) Effect of CHMP4B knock down on the secretion of IL-10 and IL-11. NIH-3T3 cells were treated with RSL3 (2 µM) for 24h. Supernatants were filtered and the cytokine composition was analyzed by the proteome profiler mouse XL cytokine Array kit. Serpin E1 was used as internal control. B) qRT-PCR quantification of transcriptional induction of IL-10 in NIH-3T3 cells treated with RSL3 (1 µM) for 2h. RPL37A was used as internal control.

## Discussion

Despite the rapid progress made in recent years on the understanding of ferroptosis, how the final and key step of plasma membrane rupture is executed remains unclear. Here we examined the cellular alterations during ferroptosis execution, including increase in cytosolic Ca^2+^, cell shape changes and lipid peroxidation, and correlated them with plasma membrane disruption that marks the final point of cell death. We found that GPX4 inactivation induces a rise in cytosolic Ca^2+^ prior to complete plasma membrane disruption. Unlike lipid peroxidation, both events are driven by osmotic forces and linked to the formation of small pores of few nanometers in the plasma membrane. Sustained increase in cytosolic Ca^2+^ correlates with the activation of the ESCRT-III machinery for membrane repair, which delays the kinetics of cell death likely by counterbalancing membrane damage upon ferroptosis. Importantly, activation of ESCRT-III during ferroptosis functions in the modulation of the inflammatory microenvironment of the dying cells, which is a common feature with other lytic types of regulated cell death such as necroptosis (Gong et al., 2017a, Gong et al., 2017b, Yoon, Kovalenko et al., 2017) and pyroptosis (Ruhl et al., 2018).

Alteration of plasma membrane permeability leading to final cell bursting is a feature of ferroptosis that is common with other mechanisms of regulated necrosis such as necroptosis and pyroptosis (Espiritu et al., 2019, Frank & Vince, 2018, Grootjans, Vanden Berghe et al., 2017, Ros et al., 2017, Sborgi et al., 2016). In necroptosis and pyroptosis, membrane permeabilization is mediated by dedicated protein cell-death effectors encoded in the genome (Chen et al., 2016, Grootjans et al., 2017, Sborgi et al., 2016). In contrast, ferroptosis is special in that membrane damage does not seem to require any specific protein machinery, but results from overwhelming accumulation of toxic lipid peroxides due to insufficient GPX4 activity (Dixon, 2017, Yang & Stockwell, 2016). How lipid peroxidation leads to plasma membrane disruption remains obscure (Agmon & Stockwell, 2017).

Here we identified that plasma membrane disruption in ferroptosis is caused by imbalance in osmotic forces due to the opening of small nanopores at the plasma membrane. In agreement with this, recent studies have suggested that accumulation of oxidized-PUFA-containing phospholipids in the membrane during ferroptosis would induce membrane thinning together with an increase in membrane curvature (Agmon, Solon et al., 2018). However, questions like the nature of the pores formed in ferroptosis and if they are transient structures composed only by lipids or whether specific proteins are involved deserve further investigation. Activation of osmotic forces leading to ion imbalance and cell bursting represents a common feature with necroptosis executed by MLKL and pyroptosis mediated by GSDMD (Ros et al., 2017, Sborgi et al., 2016). In this scenario, our results position the opening of pores at the plasma membrane as a core mechanism responsible for the execution of different types of regulated necrosis.

Different ion fluxes take place upon membrane damage, of which Ca^2+^ has a critical role linking increase in plasma membrane permeability to the activation of intracellular signaling pathways (Andrews & Corrotte, 2018, Bouillot, Reboud et al., 2018, Carafoli & Krebs, 2016, Zhivotovsky & Orrenius, 2011). In this context, rise in cytosolic Ca^2+^ has been related with the initiation and execution of different cell death processes including apoptosis, necroptosis, and pyroptosis, or unregulated necrosis (Bouillot et al., 2018, Espiritu et al., 2019, Mattson & Chan, 2003, Zhivotovsky & Orrenius, 2011). In the case of ferroptosis, contradictory observations suggested that Ca^2+^ has no impact on ferroptosis (Dixon et al., 2012, Wolpaw, Shimada et al., 2011) or linked extracellular Ca^2+^ influx and oxidative cell death with the inhibition of the *X*_*c*_ transport system by glutamate (Maher, van Leyen et al., 2018).

Here we show that Ca^2+^ fluxes can be induced at different stages upon erastin-1 or RSL3 treatment. When ferroptosis is triggered by erastin-1, there is an early increase in cytosolic Ca^2+^ event that takes place quickly after treatment but is not inhibited by Fer-1, as well as a later one that is common with the single Ca^2+^ influx event observed in RSL3-induced ferroptosis and that can be inhibited by Fer-1. Our findings could therefore reconcile possible contradictions about the essential role of Ca^2+^ in ferroptosis, since the type of fluxes that is activated downstream GPX4 inactivation strongly depends on the treatment that is used.

The cysteine/glutamate antiporter system *x*_*c*_ is the most upstream effector of the erastin-1-induced pathway. Inhibition of this system decreases the intracellular levels of cysteine, which is required for the synthesis of GSH (Bridges, Natale et al., 2012, Dixon et al., 2014). The fact that early Ca^2+^ increase activated by erastin-1 is not completely blocked by Fer-1 indicates that it is unrelated to ferroptosis. This Ca^2+^ signaling could be triggered by the inhibition by erastin-1 of the voltage-dependent anion-selective channel proteins 2 and 3 (VDAC2 and 3) (Dixon et al., 2012, Maldonado, Sheldon et al., 2013), which are involved in the regulation of cellular Ca^2+^ homeostasis (Rizzuto, De Stefani et al., 2012). In contrast, ferroptosis-specific increase in Ca^2+^ observed upon erastin-1 and RSL3 treatments is sustained and linked to membrane damage, as it was inhibited by osmotically active agents. We cannot discard the possibility that it is also a result of the indirect activation of endogenous Ca^2+^ channels, a process that is also connected to membrane damage (Ousingsawat et al., 2017, Ousingsawat, Wanitchakool et al., 2018, Simoes, Ousingsawat et al., 2018). Further work will be necessary to identify the cellular components that mediate ferroptosis-related Ca^2+^ fluxes and their role in ferroptosis progression.

Upon plasma membrane injury, local increase of cytosolic Ca^2+^ activates a number of membrane repair mechanisms that heal perforated sections of the membrane (Andrews & Corrotte, 2018, Jimenez & Perez, 2017). Recently, the ESCRT-III complex has been identified as a common machinery that mediates plasma membrane repair by shedding and trapping the damaged sections of the membranes into intracellular or extracellular vesicles in response to the increase of cytosolic Ca^2+^ triggered upon necroptosis and pyroptosis (Gong et al., 2017a, Gong et al., 2017b, Ruhl et al., 2018). As a result, the ESCRT-III machinery regulates the kinetics of necroptosis and pyroptosis and impacts on the signals released from dying cells, on intercellular communication, and on the activation of the immune system (Gong et al., 2017a, Gong et al., 2017b, Ruhl et al., 2018, Yoon et al., 2017). Here we found that the ESCRT-III machinery is also relevant to protect cells against ferroptosis. We correlated kinetically the activation of ESCRT-III with the increase in cytosolic Ca^2+^ concentration upon membrane damage in ferroptosis and showed that ESCRT-III depletion accelerates the kinetics of ferroptosis.

The lag time between membrane damage and cell bursting represents a time frame in which the cells can communicate with the microenvironment (Espiritu et al., 2019). By modulating the kinetics of cell death, the ESCRT-III machinery might thus affect the inflammatory factors that are released from ferroptotic cells, as previously proposed for necroptosis and pyroptosis (Gong et al., 2017a, Gong et al., 2017b, Ruhl et al., 2018, Yoon et al., 2017). In agreement with this, we discovered that ESCRT-III plays a role on keeping in check the levels of IL-10 secreted from ferroptotic cells. As this is a multifunctional anti-inflammatory cytokine that limits and ultimately terminates inflammatory responses, its downregulation could be crucial for the adequate activation of the immune system in the ferroptotic microenvironment. Our findings expand the role of ESCRT-III as a general machinery for counterbalancing regulated necrosis and its inflammatory consequences.

In summary, here we defined the cellular hallmarks of ferroptosis associated with plasma membrane permeabilization upon GPX4 inactivation. Activation of the ferroptotic pathway leads to sustained increase of cytosolic Ca^2+^ levels as a result of membrane damage and precedes final cell bursting. These processes are governed by osmotic pressure and result from the formation of small pores of a few nanometers in the plasma membrane. Membrane damage and the consequent activation of Ca^2+^ fluxes correlate with the activation of the ESCRT-III machinery, which counterbalances ferroptosis to modulate the kinetics of cell death. Our results that ESCRT-III activity restricts the levels of IL-10 in the ferroptotic microenvironment reveal a role of this machinery in the modulation of the inflammatory outcome of ferroptosis. These findings support a model in which plasma membrane pore formation and membrane repair emerge as central and opposite mechanisms that control the progression of different types of regulated necrosis and their inflammatory signature.

## Experimental procedures

### Reagents and antibodies

Erastin-1, RSL3, Fer-1 and Nec-1s were purchased from Biomol (Germany). PI and all PEGs were from Sigma-Aldrich (Germany). zVAD was obtained from APEXBIO (Houston, TX, USA). Fluo4-AM and C11 BODIPY 581/591 were provided by Thermofisher (Germany). CHMP4B siRNA (L-018075-01-005) was purchased from Dharmacon (Germany). The CHMP4B-eGFP and CHMP4B-mCherry constructs were kindly provided by Prof. Christian Wunder (Institut Curie, Paris, France).

### Cell lines, culture conditions and transfection

NIH-3T3 cells were obtained from Prof. Dr. Andreas Linkermann (University Hospital Carl Gustav Carus at the Technische Universität Dresden). HT-1080 and Mda 157 cells were obtained from Dr. Marcus Conrad (Helmholtz Zentrum München) under the terms of a material transfer agreement. All cells were cultured in low glucose DMEM (Sigma, Germany) supplemented with 10% FBS and 1% penicillin-streptomycin (ThermoFisher, Germany), and grown in a humidified incubator containing 5% CO_2_ at 37 °C. The cells were frequently passaged at sub-confluence, and seeded at a density of 0.5–5 × 10^4^ cells/mL. For knock down experiments, cells were transfected with Lipofectamine 2000 (Invitrogen, Germany) with specific siRNA, for 48 hours in 6-well (western blot) or 12-well (FACS) plates.

### Flow cytometry measurements

The plasma membrane integrity was tested by flow cytometry measuring the ability of cells to exclude PI. Flow-cytometric analyses were conducted using CytoFlex and data analysed using the FACSDiva software (Beckman Counter, Germany). After treatment, both attached and non-attached cell populations were collected. Cells were washed twice with cold PBS, centrifuged (2500 *g*, 5 min, 4°C), and resuspended in PBS containing Fluo4-AM (1 µM). After 30 min of incubation at 37 °C, cells were washed, resuspended in PBS containing PI (2 µg/mL) and a total of 10 000 cells were analyzed by flow cytometry.

### Bright field and confocal microscopy

Cells were seeded in DMEM in IBIDI 8-well chambers (Ibidi, Germany) 24 hours before experiment. The day after, cells were washed with PBS to replace the media by phenol red free DMEM (Sigma-Aldrich, Germany) supplemented with FBS and antibiotics. Cells were loaded with 1 µM Fluo-4 AM, 2 µg/mL PI or 1 µM of C11 BODIPY 581/591 for 30 min at 37°C. All images were acquired with a Zeiss LSM 710 ConfoCor3 microscope (Carl Zeiss, Jena, Germany) equipped with incubator at 37°C and 5% CO_2_. Time lapse imaging with z-stack acquisition was carried out before and after ferroptosis induction. Transmitted light and fluorescence images were acquired through a Zeiss C-Apochromat 40X, NA = 1.2 water immersion objective onto the sample. Excitation light came from argon ion (488 nm) or HeNe (561 nm) lasers. All images were processed in Fiji.

### Lipid peroxidation measurements

The oxidation ratio of C11 BODIPY 581/591 was calculated as an indicator of lipid peroxidation per cell following the equation:

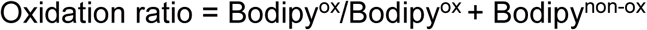

Where Bodipy^non-ox^ corresponds to the non-oxidized fraction of the probe and was estimated based on the fluorescence intensity per pixel from the red and Bodipy^ox^ corresponds to the oxidized fraction and was calculated based on the fluorescence intensity per pixel from the green channel fluorescence images.

### Cytokine Array

Culture supernatants from NIH-3T3 cells treated with RSL3 for 24 hours were analyzed for secreted cytokines/chemokines using Proteome Profiler Mouse XL Cytokine Array (R&D Systems) according to the manufacturer’s instructions. Signal intensities of the individual cytokines/chemokines were quantified by densitometry using the software ImageJ and normalized to the non-treated controls.

### RNA isolation, cDNA generation and quantitative PCR

NIH-3T3 cells were treated with RSL3 for 2 hours, and RNA was isolated using the RNeasy Mini Kit (Qiagen). Subsequently, cDNA was generated by reverse transcription with the cDNA Synthesis Kit (Biozym) and random hexamer primers. Resulting cDNA was used for qPCR which was performed on a LightCycler 480 II system (Roche) using GreenMasterMix (Genaxxon). Relative gene expression was calculated with the 2 ΔΔCt method (Livak and Schmittgen, 2001). HPRT served as a reference gene. The following primers were used for qPCR:

HPRT 5’-TCAGTCAACGGGGGACATAAA-3’ and

5’-GGGGCTGTACTGCTTAACCAG-3’;

IL10 5’-GCTCTTACTGACTGGCATGAG-3’ and

5’-CGCAGCTCTAGGAGCATGTG-3’;

IL11 5’-TGTTCTCCTAACCCGATCCCT-3’ and

5’-CAGGAAGCTGCAAAGATCCCA-3’.

### Statistical methods

All measurements were performed at least 3 times and results are presented as mean ± standard deviation.

## Author contributions

L.P., R.A.E. and U.R. performed confocal and flow cytometry experiments and analyzed data. U.R., A.S. and S.H. designed and conducted the cytokine experiments. All authors contributed to experimental design and manuscript writing. A.J.G.S conceived the project and supervised research.

## Acknowledgments

L.P., R.A.E., and U.R. acknowledge the Alexander von Humboldt Foundation for supporting their postdoctoral research at the Interfaculty Institute of Biochemistry, Eberhard-Karls-Universität Tübingen, Germany. Current work of U.R. is supported by Eberhard-Karls-Universität Tübingen. R.A.E would also like to express his gratitude to De La Salle University (Manila, Philippines) for additional financial support. This work was partially supported by the DFG grant AG1641/6-1.

## Main figures and legends

**Figure S1:**
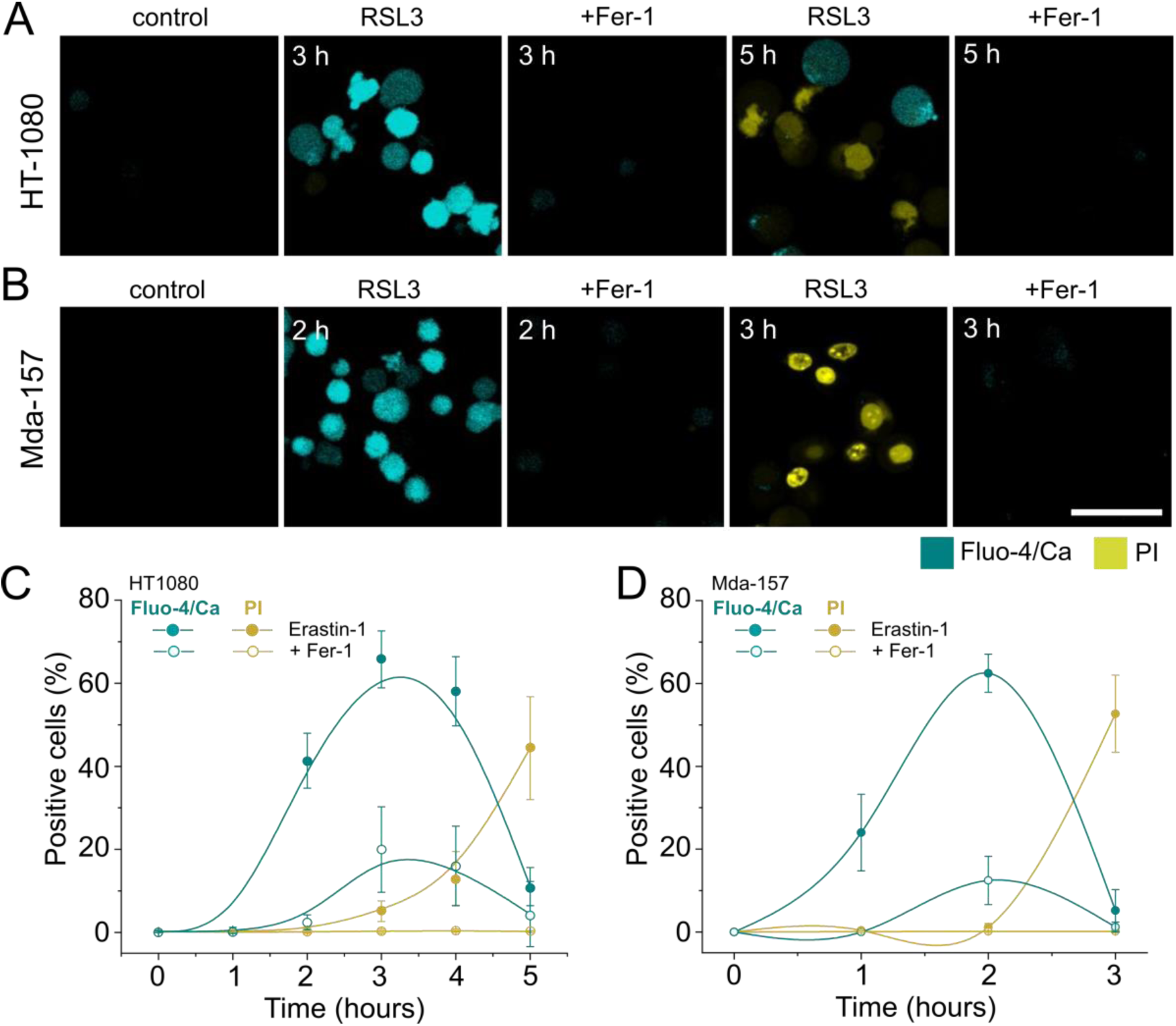
Level of cytosolic calcium increase and cell death during ferroptosis in RSL3-treated cells. A and B) Representative confocal images of cells treated with RSL3 in the presence or not of Fer-1. Scale bar, 50 µm. C and D) Kinetics of increase of cytosolic Ca^2+^ and plasma membrane breakdown in cells treated with RSL3 in the presence or not of Fer-1. The values represent the mean and the standard deviation of at least three independent experiments.

**Figure S2:**
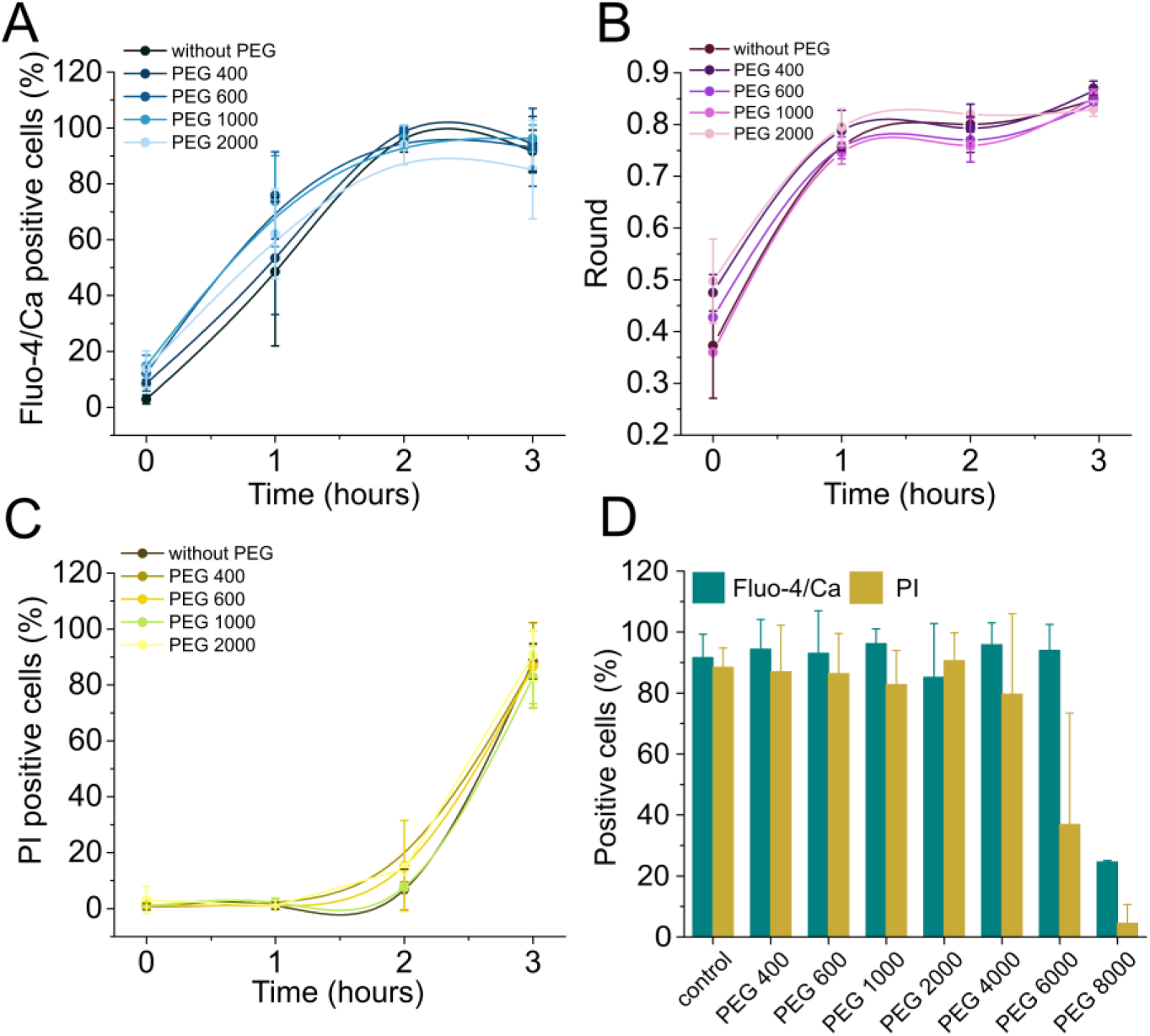
PEGs of smaller sizes did not provide osmotic protection against ferroptosis. A) Kinetics of increase of cytosolic Ca^2+^, B) change in cell shape, and C) PI intake in NIH-3T3 cells treated with RSL3, in the presence or not of PEGs of different sizes. D) Ca^2+^ signal and PI intake in NIH-3T3 cells treated 3 hours with RSL3 with or without PEGs of different sizes. Each data point represents the mean of at least six replicas from three independent experiments. At least 100 cells were analyzed for each replica. Concentrations: RSL3 (2 µM), PEGs (5 mM). PEG sizes: 400 (0.56 nm), 600 (0.69 nm), 1000 (0.94 nm), 2000 (1.6 nm), 4000 (1.8 nm), 6000 (2.3 nm) and 8000 (2.7 nm).

**Figure S3:**
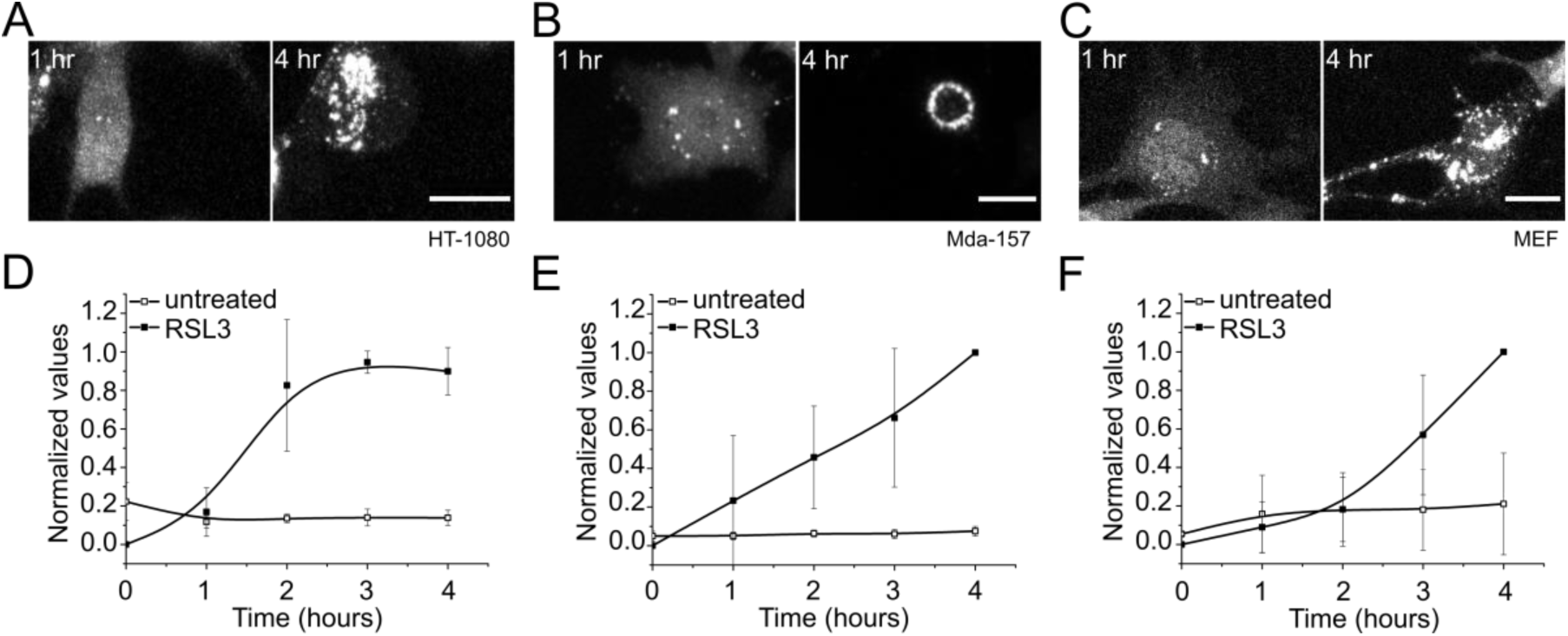
CHMP4B activation in RSL3-treated cells to protect from cell death via membrane repair. A-C) Representative images of HT-1080, Mda-157, and MEF cells transiently transfected with CHMP4B-eGFP and treated with RSL3, and monitored for CHMP4B punctae formation. Scale bar, 20 µm. D-F) Kinetics of appearance of CHMP4B punctae in the corresponding cells, treated or not with RSL3. (N = 3 – 8 cells).

